# Immunization of mice with chimeric antigens displaying selected epitopes confers protection against intestinal colonization and renal damage caused by Shiga toxin-producing *Escherichia coli*

**DOI:** 10.1101/783829

**Authors:** David A. Montero, Felipe Del Canto, Juan C. Salazar, Sandra Cespedes, Leandro Cádiz, Mauricio Arenas-Salinas, José Reyes, Ángel Oñate, Roberto M. Vidal

**Author notes:** Corresponding author. Correspondence should be addressed to R.V.

## Abstract

Shiga toxin-producing *Escherichia coli* (STEC) cause diarrhea and dysentery, which may progress to hemolytic uremic syndrome (HUS). Vaccination has been proposed as a preventive approach against STEC infection; however, there is no vaccine for humans and those used in animals reduce but do not eliminate the intestinal colonization of STEC. The OmpT, Cah and Hes proteins are widely distributed among clinical STEC strains and are recognized by serum IgG and IgA in patients with HUS. Here, we develop a vaccine formulation based on two chimeric antigens containing epitopes of OmpT, Cah and Hes proteins against STEC strains. Intramuscular and intranasal immunization of mice with these chimeric antigens elicited systemic and local long-lasting humoral responses. However, the class of antibodies generated was dependent on the adjuvant and the route of administration. Moreover, while intramuscular immunization with the combination of the chimeric antigens conferred protection against colonization by STEC O157:H7 and the intranasal conferred protection against renal damage caused by STEC O91:H21. This pre-clinical study supports the potential use of this formulation based on recombinant chimeric proteins as a preventive strategy against STEC infections.

## Introduction

Shiga toxin-producing *Escherichia coli* (STEC) are a group of food-borne pathogens causing acute and bloody diarrhea, which may progress to life-threatening complications such as hemolytic uremic syndrome (HUS).^1^ To date there is no specific treatment for STEC infection and antibiotic use is contraindicated due to increased risk of HUS development.^2^ However, some drugs have been specifically designed to protect against the effects of the presence of Shiga toxins and are in different stages of clinical trials.^3,4^ While STEC O157:H7 is the serotype most frequently associated with diarrhea outbreaks and HUS cases worldwide, there are other serotypes, the incidence and impact of which on public health and the food industry have increased.^5,6^

STEC colonizes the human colon and produces Shiga toxins (Stx) that can enter the blood stream and disseminate to organs such as the kidneys and central nervous system. Once Stx reach the target organs and enter the cells, the toxins inhibit protein synthesis, leading to autophagy and apoptosis and ultimately tissue damage, which may lead to HUS.^7^

To colonize the human colon, STEC requires several virulence factors like those encoded in the locus of enterocyte effacement (LEE) pathogenicity island (PAI). LEE-mediated adherence causes the formation of the “attaching and effacing” lesion and loss of microvilli of the intestinal epithelial cells.^8^ In addition, STEC strains lacking LEE (LEE-negative STEC) harbor other PAIs like the Locus of Adhesion and Autoaggregation (LAA), which encodes virulence factors involved in intestinal colonization.^9,10^ In fact, the presence of two or more PAIs in single isolates of clinically relevant STEC serotypes is common, suggesting that the cumulative acquisition of mobile genetic elements encoding virulence factors may contribute additively or synergistically to pathogenicity.^10,11^

Vaccination of the infant population, which is the highest-risk group for STEC infections, and animal reservoirs have been proposed as a preventive approach that could reduce their incidence and prevalence. However, there is no approved STEC vaccine for humans, and commercial vaccines used in cattle reduce but do not eliminate colonization and shedding of these bacteria.^12^ Therefore, the development of an effective STEC vaccine is still underway. STEC proteins involved in attachment to host tissues are eligible targets for vaccine development, as they determine initial steps during infection; however, the selection of antigens that may provide a broadly and protective immune response among their diverse adhesion and colonization mechanisms is a pivotal point to consider.^13^ An additional difficulty for the development of an effective STEC vaccine has been the lack of an animal model of infection that can reproduce the pathologies caused in humans.^14^ Despite these limitations, several STEC vaccine candidates have been evaluated in laboratory animals (mice, rats and rabbits) and in cattle, with promising results. They include Stx subunit-based vaccines,^15–17^ protein and peptide-based vaccines,^17–21^ attenuated bacteria-based vaccines,^22^ bacterial ghost-based vaccines,^23^ DNA-based vaccines,^24,25^ and more recently nanoparticle-based vaccines.^26^ While most of these vaccine candidates are based on LEE-encoded antigens and Stx subunits, there are several antigens encoded outside LEE that are expressed *in vivo* during human infection that could be suitable targets for vaccine development.^13,27^

For instance, <u>O</u>uter <u>m</u>embrane <u>p</u>rotease T (OmpT) and <u>C</u>alcium binding antigen 43 <u>h</u>omologue (Cah) proteins have been shown to be recognized by IgG and IgA antibodies present in sera from patients who develop HUS (hereinafter referred to as HUS sera). Notably, the *ompT* gene has been identified in almost all clinical STEC strains and, in the case of the *cah* gene, its detection frequency is higher than 70%.^13^ Another promising antigen is the <u>H</u>emagglutinin from <u>S</u>TEC (Hes), which is recognized by IgG present in HUS sera.^13^ In addition, the *hes* gene, which is carried by the LAA PAI, is identified in about 40% and 46% of LEE-negative STEC strains isolated from humans and cattle, respectively.^9,10,28^ Thus, these three antigens are widespread among clinical STEC strains, but it is also important to note that they are mostly absent in commensal *E. coli* strains,^9,10,13^ which could diminish the probability of cross-reactivity with commensal microbiota. Nevertheless, the production and purification of outer membrane proteins (OMPs) such as OmpT, Cah and Hes poses a challenge due to their partially hydrophobic surfaces, flexibility and lack of stability that affect their solubility and efficient purification. In addition, strong detergents are used in the purification of this class of proteins and therefore the loss of conformational epitopes may impair their antigenicity and efficiency as immunogens.^29^

To circumvent these issues and to develop a STEC vaccine targeting the OmpT, Cah and Hes proteins, we implemented a vaccine development approach based on the identification of linear B-cell epitopes for the design of chimeric antigens that include them. Here we demonstrate that the immunization of mice with chimeric antigens displaying selected epitopes of OmpT, Cah and Hes proteins induces immune responses that reduce intestinal colonization and prevent renal damage caused by STEC. Overall, this study revealed the feasibility of using such type of formulation based on recombinant chimeric proteins against STEC colonization and more relevantly to protect against kidney damage by Shiga toxin-producing *Escherichia coli*.

We anticipate that our comprehensive experimental approach will contribute to the development and evaluation of future chimeric antigen-based vaccines, and while our candidate was initially intended to protect humans against colonization/infection by STEC, we believe that it could also have an effect on STEC elimination in the animal reservoir (bovine and pig) and realistically such a trial is most likely to be conducted first.

## Results

### The OmpT, Cah and Hes proteins have several linear B-cell epitopes that are recognized by IgG and IgA from HUS sera but not from control sera

An overview of our vaccine development approach is shown in Figure 1. We carried out a high-throughput screening of linear B-cell epitopes in the OmpT, Cah and Hes proteins by using a peptide microarray assay (see Methods). A total of 6, 6 and 5 epitope-like spot patterns were identified in the peptide slides of OmpT, Cah and Hes proteins (**Fig. 2a-2c**), respectively. As a complementary approach, we used a number of immunoinformatics tools and found that 13 out of 17 of the experimentally identified B-cell epitopes were predicted *in silico* by one or more algorithms (**Table 1**). This result highlights the advances in the accuracy of these bioinformatics tools. On the other hand, it is known that MHC class II (MHC-II) epitopes included in peptide vaccines enhance T-cell-dependent antibody responses ^30^. Thus, we also performed an *in silico* analysis for the prediction of MHC-II binding peptides and found two putative T-cell epitopes in the OmpT and Cah proteins (**Table 2**).

**Fig. 1.**
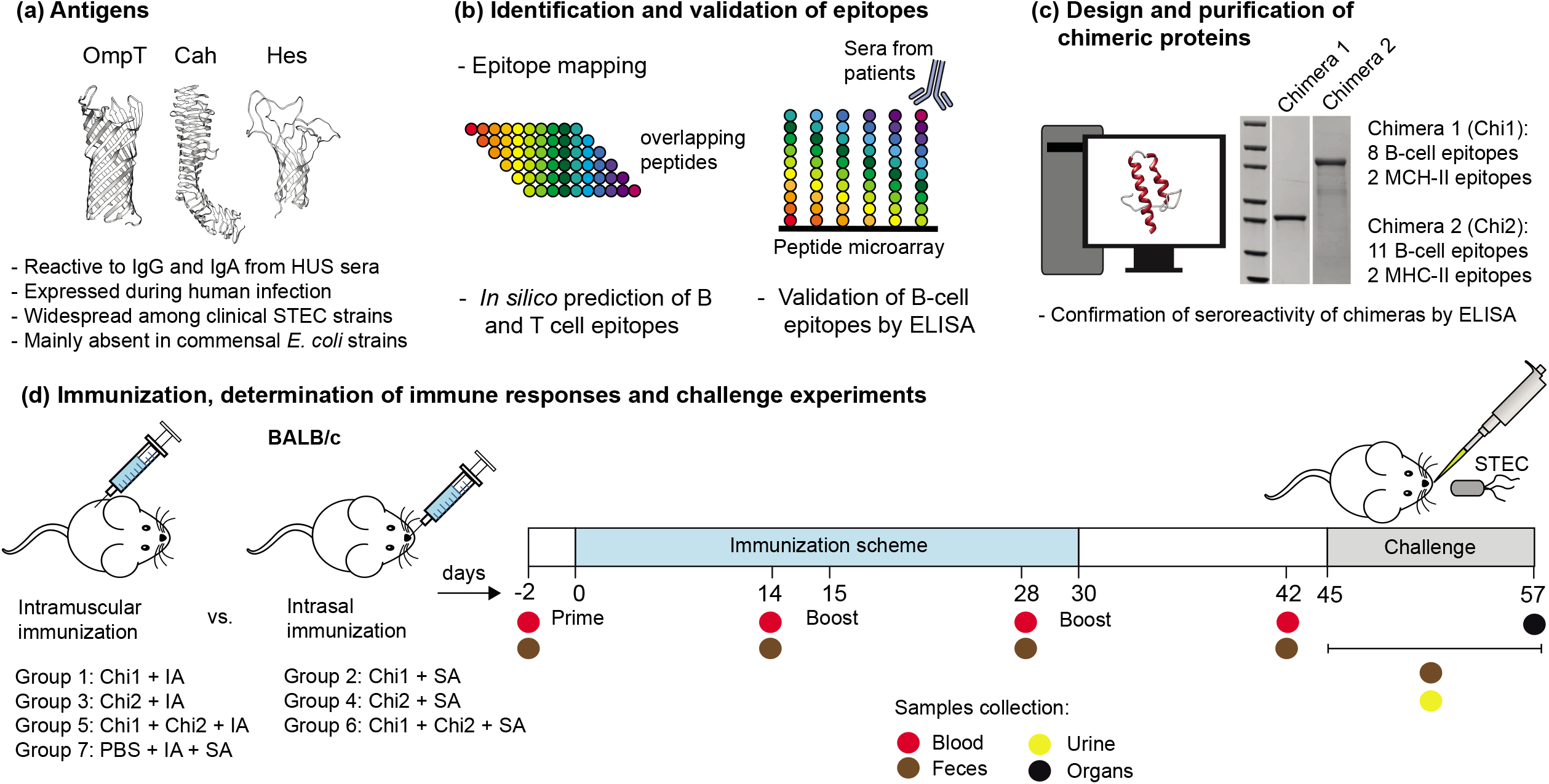
Experimental design for the development and evaluation of the chimeri-cbased STEC vaccine. **a**) We selected the OmpT, Cah and Hes proteins as suitable targets against STEC based on their antigenic properties, frequency of detection among clinical STEC strains and absence among commensal *E. coli* strains. **b**) The proteins were analyzed by immunoinformatics tools and a peptide microarray assay for B-cell epitope prediction and mapping, respectively. MHC-II epitopes were also predicted *in silico*. Several B-cell epitopes were validated by ELISA using a collection of HUS sera. **c**) Selected epitopes were used to design two chimeric proteins that were expressed and purified. The reactivity of the chimeric proteins to IgG and IgA present in HUS sera was also confirmed. **d**) BALB/c mice were immunized with different vaccine formulations by intramuscular or intranasal route using Imject Alum (IA) or Sigma adjuvant (SA), respectively. Systemic and local humoral responses were subsequently determined. The protection conferred by immunizations was evaluated in the streptomycin-treated mouse model by challenge with STEC O157:H7 and O91:H21 strains. Bacterial shedding and intestinal colonization were determined for the STEC O157:H7 infected mice. Renal damage was examined in the STEC O91:H21 infected mice.

**Fig. 2.**
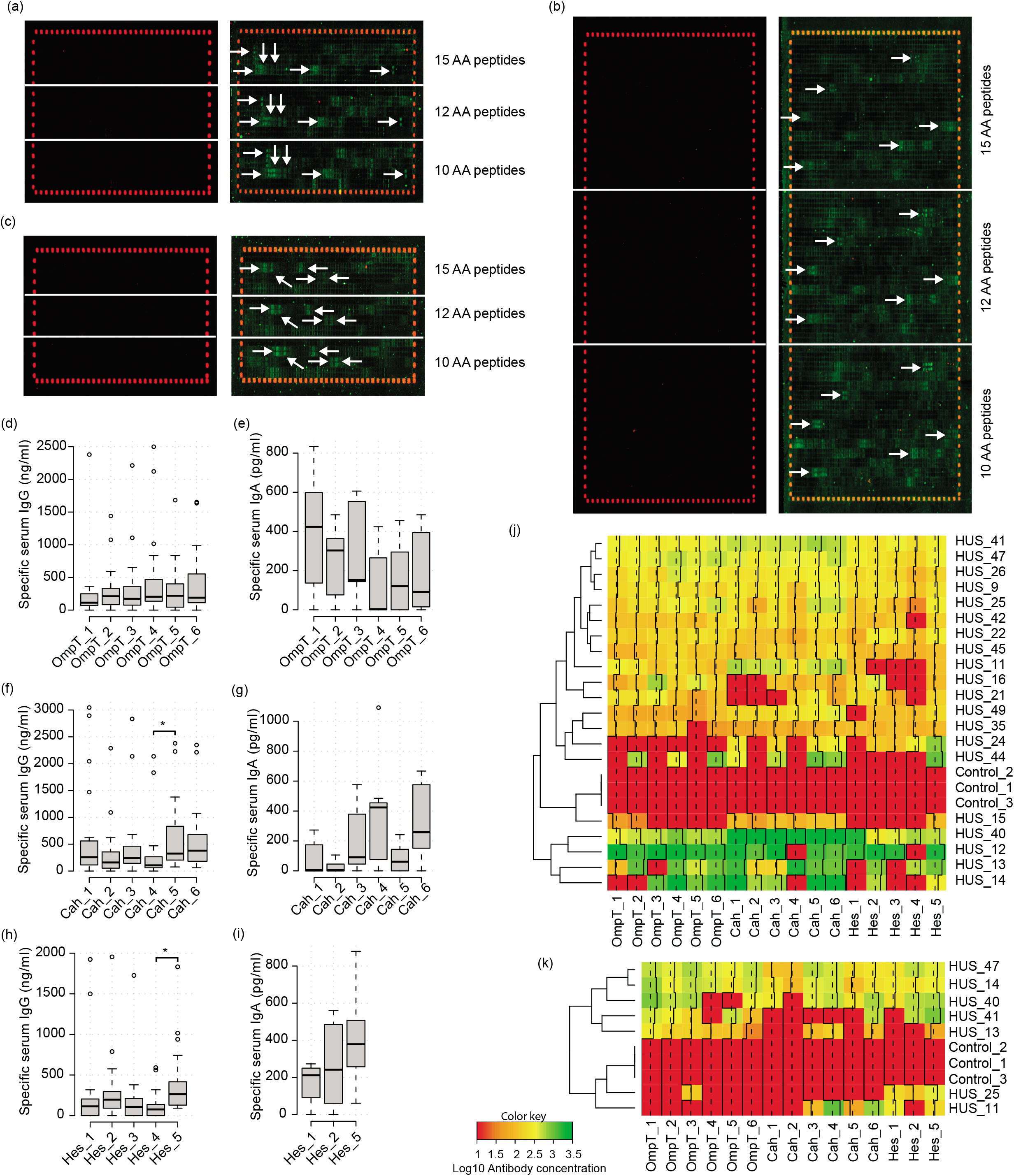
Identification and validation of linear B-cell epitopes of the OmpT, Cah and Hes proteins. **a, b and c)** Peptide microarray assays. Peptide slides containing 15, 12 and 10 AA peptides derived from de OmpT (a), Cah (b) and Hes (c) proteins were incubated with a mix of three HUS sera at a dilution of 1:100. After washing, staining was performed with secondary anti-human IgA DyLight800 antibody at a dilution of 1:1000. Control peptides (red spots) framing the peptide slides were staining with specific monoclonal DyLight680 antibody at a dilution of 1:2000. Control peptide slides incubated with anti-human IgA DyLight800 antibody and the specific monoclonal DyLight680 antibody are shown in the left panels. Epitope-like spot patterns are indicated by white arrows. **d-i**) Tukey box plots showing concentrations of IgG and IgA present in individual HUS sera (n=20) that are reactive to short peptides containing B-cell epitopes of the OmpT, Cah and Hes proteins. A lower number of HUS sera and Hes epitopes were assessed due to sera availability. Tukey box plots show the 25^th^ to 75^th^ percentiles, with the median indicated by the horizontal line inside the box. Data analysis was by Kruskal-Wallis test, followed by Dunn’s multiple comparison test. *P < 0.0 5 was considered significant. **j** and **k**) Heatmaps show the logarithm of the IgG and IgA concentration of each serum (HUS or control sera) that recognizes a specific short peptide, respectively. Data were clustered hierarchically using Euclidean distance and complete linkage analyses. Each row represents a different serum and each column a specific epitope. The average (dotted line) and histogram (solid line) of the values obtained by each peptide are indicated in the columns. The color key indicates the value of the logarithm of the antibody concentration. The figure was made using the “gplots” ^90^ package in R ^91^.

**Table 1.** Linear B-cell epitopes identified in the OmpT, Cah and Hes proteins

| Protein | Epitope name | Epitope Mapping | *In silico* B-cell prediction tools |  |  | Reactivity to HUS sera by ELISA. No (%) |  | Reactivity to control sera by ELISA. No (%) |  |
| --- | --- | --- | --- | --- | --- | --- | --- | --- | --- |
|  |  |  | BepiPred 2.0 | Kolaskar and Tongaonker | Ellipro | IgG | IgA | IgG | IgA |
| **OmpT** | OmpT_1 | Yes | Yes | No | Yes | 17/20 (85) | 5/7 (71) | 0/3 (0) | 0/3 (0) |
|  | OmpT_2 | Yes | Yes | No | Yes | 18/20 (90) | 5/7 (71) | 0/3 (0) | 0/3 (0) |
|  | OmpT_3 | Yes | Yes | Yes | Yes | 16/20 (80) | 6/7 (86) | 0/3 (0) | 0/3 (0) |
|  | OmpT_4 | Yes | Yes | Yes | Yes | 18/20 (90) | 3/7 (43) | 0/3 (0) | 0/3 (0) |
|  | OmpT_5 | Yes | Yes | No | Yes | 16/20 (80) | 4/7 (57) | 0/3 (0) | 0/3 (0) |
|  | OmpT_6 | Yes | Yes | No | Yes | 18/20 (90) | 5/7 (71) | 0/3 (0) | 0/3 (0) |
| **Cah** | Cah_1 | Yes | Yes | Yes | Yes | 18/20 (90) | 3/7 (43) | 0/3 (0) | 0/3 (0) |
|  | Cah_2 | Yes | No | No | No | 16/20 (80) | 2/7 (29) | 0/3 (0) | 0/3 (0) |
|  | Cah_3 | Yes | Yes | No | Yes | 19/20 (95) | 5/7 (71) | 0/3 (0) | 0/3 (0) |
|  | Cah_4 | Yes | No | No | No | 16/20 (80) | 5/7 (71) | 0/3 (0) | 0/3 (0) |
|  | Cah_5 | Yes | No | No | No | 20/20 (100) | 4/7 (57) | 0/3 (0) | 0/3 (0) |
|  | Cah_6 | Yes | No | No | No | 20/20 (100) | 6/7 (86) | 0/3 (0) | 0/3 (0) |
| **Hes** | Hes_1 | Yes | Yes | No | Yes | 14/20 (70) | 5/7 (71) | 0/3 (0) | 0/3 (0) |
|  | Hes_2 | Yes | Yes | Yes | Yes | 17/20 (85) | 5/7 (71) | 0/3 (0) | 0/3 (0) |
|  | Hes_3 | Yes | Yes | No | Yes | 14/20 (70) | N.E. | 0/3 (0) | N.E. |
|  | Hes_4 | Yes | No | No | Yes | 12/20 (60) | N.E. | 0/3 (0) | N.E. |
|  | Hes_5 | Yes | Yes | No | Yes | 20/20 (100) | 7/7 (100) | 0/3 (0) | 0/3 (0) |

N.E. Not evaluated

**Table 2.** Epitopes and chemical and physical properties of the chimeric proteins

| Protein | B-cell epitopes | Predicted T-cell epitopes | MW (kDa) | Theoretical pI | Solubility <sup>1</sup> | Estimated half-life <sup>2</sup> | Instability index <sup>3</sup> |
| --- | --- | --- | --- | --- | --- | --- | --- |
| Chimera 1 | OmpT_1, OmpT_2, OmpT_3, OmpT_4, Hes_1, Hes_2, Hes_3 and Hes_5 | Two of OmpT | 29.6 | 4.51 | 0.693 | >10 hours | 21.30 |
| Chimera 2 | OmpT_1, OmpT_6, Cah_1, Cah_2, Cah_3, Cah_4, Cah_5, Cah_6, Hes_1, Hes_2 and Hes_5 | Two of Cah | 56.7 | 4.52 | 0.666 | >10 hours | 8.02 |

<sup>1</sup> Predicted by Protein-Sol tool <sup>69</sup>. Scaled solubility value (0-1). A value greater than 0.45 predicts that the protein is soluble.
<sup>2</sup> Prediction of the time it takes for half of the amount of protein in *E. coli* to disappear after its synthesis. <sup>3</sup> The instability index provides an estimate of the stability of a protein in a test tube. A protein whose instability index is smaller than 40 is predicted as stable <sup>77</sup>.

Because the peptide microarray assay was performed with a mix of three HUS sera, we sought to confirm the reactivity of these epitopes using a larger number of sera. We also tested the reactivity of the epitopes against three sera obtained from children with no medical record of STEC-related disease (hereinafter referred to as control sera). For this, short peptides ranging from 15 to 24 amino acids (aa) which include each epitope were chemically synthesized, and their reactivity to IgG and IgA was assessed by ELISA. In general, the peptides were recognized at higher levels by IgG than by IgA from HUS sera (**Fig. 2d-i**). Furthermore, the peptides derived from the same antigen were recognized by similar levels of IgG or IgA, with the exception of Cah_5 and Hes_5, which showed a higher level of reactivity to IgG compared to Cah_4 and Hes_4, respectively (**Fig. 2f and 2h**). The frequency of reactivity of the peptides to IgG ranged from 60% (Hes_4) to 100% (Cah_5, Cah_6 and Hes_5), while for IgA it was between 29% (Cah_2) and 100% (Hes_5) (**Table 1**). As expected, a polyclonal antibody response was observed within individuals, evidenced by the variable concentration of IgG and IgA antibodies that recognize each tested peptide (**Fig. 2j-k**). Importantly, none of the peptides was recognized by IgG or IgA from control sera (**Table 1, Fig. 2j-k**). Taken together, these results indicate that the identified linear B-cell epitopes are broadly recognized and immunodominant during immune responses against STEC.

### Chimeric proteins displaying linear B-cell epitopes of OmpT, Cah and Hes proteins are recognized by IgG and IgA from HUS sera but not from control sera

We consider that the best epitopes for vaccine development are those that are conserved, broadly distributed among clinical isolates, with higher levels of immunoreactivity and surface exposition in the native antigen. As a result, the Hes_4 (lower reactivity and detection frequency) and OmpT_5 (limited surface exposition) epitopes were discarded and not used in further assays. For the *in silico* design of proteins we implemented two different approaches. Firstly, we noticed that the linear B-cell epitopes of OmpT and Hes are overlapped or consecutively arranged along the protein (**Fig. 2a and 2c**), suggesting that they form antigenic domains (AD). We took advantage of this and designed a chimeric protein containing these AD through the fusion of 135 AA and 127 AA from OmpT and Hes proteins, respectively (**Fig. 3a**). We named this protein Chimera 1 (Chi1; 262 AA and 29 kDa), which includes a total of eight B-cell epitopes (OmpT_1, OmpT_2, OmpT_3, OmpT_4, Hes_1, Hes_2, Hes_3 and Hes_5) and two predicted T-cell epitopes (**Table 2**). In the second approach, we used the passenger domain of Cah (αCah) as a carrier (keeping its epitopes) and incorporated several B-cell epitopes of OmpT and Hes (**Table 2, Fig. 3d**). Thus, this second protein named Chimera 2 (Chi2; 559 AA and 56 kDa) includes a total of eleven B-cell epitopes (OmpT_1, OmpT_6, Hes_1, Hes_2 and Hes_5, Cah_1, Cah_2, Cah_3, Cah_4, Cah_5, Cah_6) and two predicted T-cell epitopes (**Table 2**).

**Figure 3.**
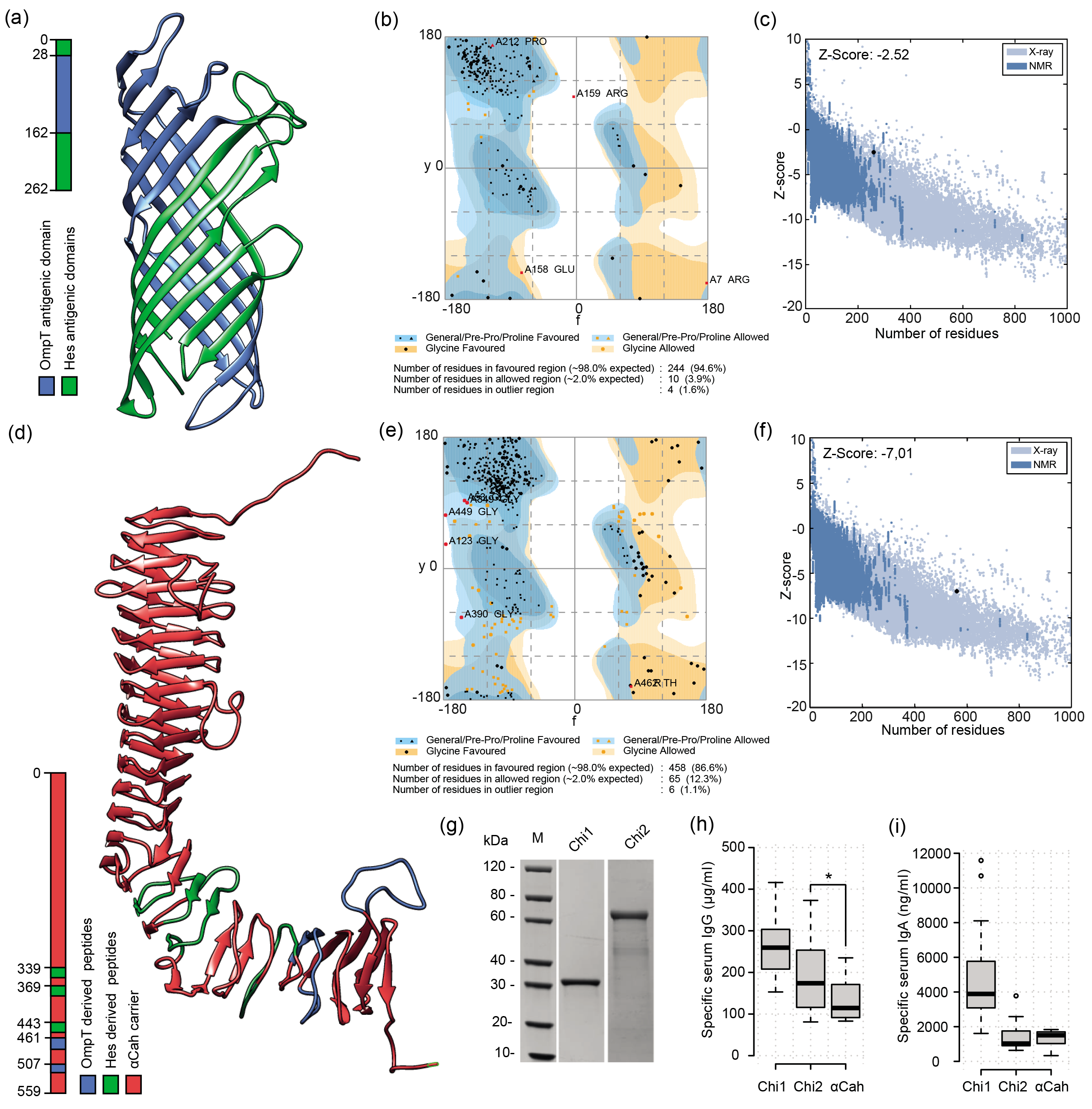
Design and production of the chimeric antigens. **a)** Predicted 3D structure of the Chimera 1 (Chi1) antigen. OmpT- and Hes-derived peptides are shown as indicated in the legend at the left. **b and c**) Quality validation of modeled Chi1 structure. Ramachandran plot (b) shows that 94.6% residues were in favored regions. Z-score plot (c) for Chi1 model obtained by ProSA-web.^81^ Dark blue and light blue regions represent z-scores of native protein structures determined by NMR and X-ray, respectively. Black spot shows z-score for the Chi1 model. **d**) Predicted 3D structure of the Chimera 2 (Chi2) antigen. OmpT- and Hes-derived peptides are shown as indicated in the legend at the left. **e and f**) Quality validation of modeled Chi2 structure. Ramachandran plot (e) shows that 86.6% residues were in favored regions. Z-score plot (f) for Chi2 model. **g**) SDS-PAGE of purified Chi1 and Chi2 proteins. M, Molecular weight standard. **h-i**) Tukey box plots showing concentrations of IgG (h) and IgA (i) present in individual HUS sera (n=20) that are reactive to Chi1, Chi2 and αCah proteins. Tukey box plots show the 25^th^ to 75^th^ percentiles, with the median indicated by the horizontal line inside the box. Data analysis was by Kruskal-Wallis test, followed by Dunn’s multiple comparison test. *P < 0.0 5 was considered significant.

We predicted the 3D structures for both chimeric proteins, which were refined and validated (see Methods) (**Fig. 3a and 3d**). A Ramachandran plot analysis revealed that 94.6% and 86.6% of amino acid residues from the Chi1 and Chi2 modeled structures were in favored regions, respectively (**Fig. 3b and 3e**). In addition, the Z-score of the Chi1 and Chi2 modeled structures were −2.52 and −7.01 respectively, which are within the range of scores found for native proteins of similar size (**Fig. 3c and 3f**). The predicted solubility, *in vivo* half-life and instability index of both chimeric proteins suggested that their expression and purification could be feasible (**Table 2**). Consistent with the above, the production of these recombinant proteins in *E. coli* showed that they are stable, water-soluble and have the predicted molecular weight (**Fig. 3g**). Further, we confirm that both the Chi1 and Chi2 proteins are recognized by IgG and IgA of HUS sera (**Fig. 3h and 3i**). We also found that the reactivity of Chi2 to IgG of HUS sera was significantly higher than αCah (**Fig. 3h**), indicating that the incorporation of B-cell epitopes of OmpT and Hes increased the seroreactivity. However, this difference was not observed in the reactivity to IgA of HUS sera (**Fig. 3i**). Importantly, none of the proteins was seroreactive to IgG and IgA of control sera (not shown).

### Chi1 and Chi2 antigens, administered alone or in combination, trigger long-lasting systemic and local humoral responses in mice

Having established that the Chi1 and Chi2 proteins are seroreactive to HUS sera, we sought to evaluate them as immunogens. For this, BALB/c mice were immunized following the scheme described in Figure 1d. Immunity achieved by vaccination is influenced to a large extent by the administration route and the type of adjuvant^31–33^. Therefore, we also compared systemic and local immune responses triggered by the chimeric antigens when administered with Imject Alum or Sigma adjuvants by intramuscular or intranasal route, respectively.

The measurement of Chi1 and Chi2-specific IgG antibodies in serum showed that mice immunized with Chi2 or Chi1 plus Chi2 by either intramuscular or intranasal route elicited significantly higher levels of IgG on days 28 and 42 than the PBS control group (**Fig. 4a**). In general, intramuscular immunization induced higher levels of specific IgG antibodies than the intranasal route, this difference being significant in mice immunized with Chi1 plus Chi2. Similarly, specific IgA antibodies in serum were significantly higher in mice immunized with Chi2 or Chi1 plus Chi2 by both administration routes on days 14, 28 and 42 than in the PBS control group (**Fig. 4b**). It was also observed that intramuscular immunization with Chi1 plus Chi2 induced higher levels of specific IgA antibodies than the intranasal route, this difference being significant on day 42. In contrast, there were no differences on the levels of specific IgG and IgA antibodies in serum between mice immunized with Chi1 and the PBS control group (**Fig. 4a-b**). Regarding specific IgM antibodies in serum, all vaccine formulations and administration routes induced significant higher antibody levels on days 14, 28 and 42 than the PBS control group (**Fig. 4c**).

**Figure 4.**
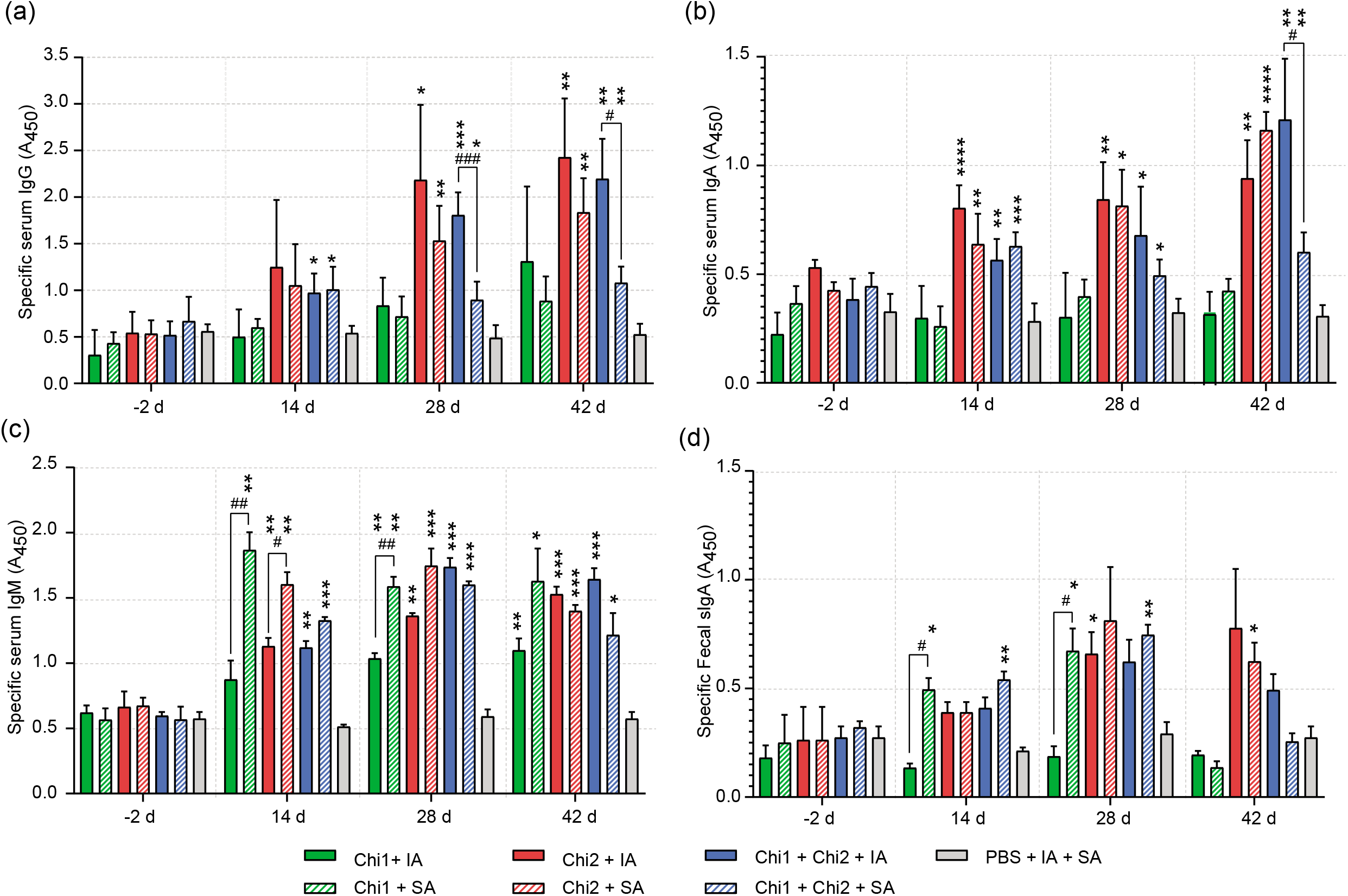
Humoral immune responses triggered by immunization with the chimeric antigens. Sera obtained at days −2, 14, 28 and 42 post-immunization were diluted 1:100 and used for the determination of specific IgG (a), IgA (b) and IgM (c) by ELISA. Fecal sIgA (d) was also determined from feces collected on days −2, 14, 28 and 42. The ELISA plates were coated with 100 μl of each antigen (Chi1, Chi2 and an equimolar mix Chi1+Chi2) at a final concentration of 1 μg/ml in carbonate-bicarbonate buffer (pH 9.6) and incubated overnight at 4 °C. Then, the plates were blocked, washed and incubated with different dilutions of each serum. The results are expressed as means ± SD of absorbance values at 450 nm (A_50_), which were obtained from individual sera or fecal suspensions of five mice per group. Experimental groups are shown as indicated by legend at the bottom. Data analysis was by a two-way ANOVA, followed by Tukey’s multiple comparison test. P < 0.0 5 was considered significant. Asterisks (*) indicate significant differences between the immunized mice and the PBS control group. Number signs (#) indicate significant differences between administration routes.

To evaluate the induction of mucosal responses, specific secretory IgA (sIgA) antibodies were determined in feces. Mice immunized with Chi1 and Chi1 plus Chi2 by intranasal route elicited significantly higher levels of specific sIgA on days 14 and 28 than the PBS control group (**Fig. 4d**). However, the sIgA levels for both experimental groups were similar on day 42 compared to the PBS control group. On day 42, only mice immunized with Chi2 by intranasal route elicited significantly higher levels of specific sIgA than the PBS control group. Taken together, these results indicate that immunization with Chi1, Chi2 or Chi1 plus Chi2 induces a systemic and local humoral response influenced by the type of adjuvant and the administration route as long as 42 days after the third immunization.

### Chi1 and Chi2 antigens administered in combination by intramuscular route reduce intestinal colonization and fecal shedding of STEC O157:H7

We next evaluated whether immune responses elicited by vaccination with the chimeric antigens may confer protection against intestinal colonization by STEC. For this, two weeks after the last booster immunization, mice were treated with streptomycin and then orally challenged with STEC O157:H7 str. 86-24 (**Fig. 1d;** see Methods). Notably, mice immunized with Chi1 plus Chi2 by intramuscular route showed a significantly lower fecal shedding of the STEC 86-24 strain from day 8 post-infection to the end of the experiment (day 12) compared to the PBS control group (**Fig. 5a**). In mice immunized by the intranasal route, only the Chi2 group showed a slight but significant decrease in fecal shedding of STEC 86-24 strain on days 7 and 8 compared to the PBS control group. However, on day 9 and later, this difference was not observed (**Fig. 5b**).

**Figure 5.**
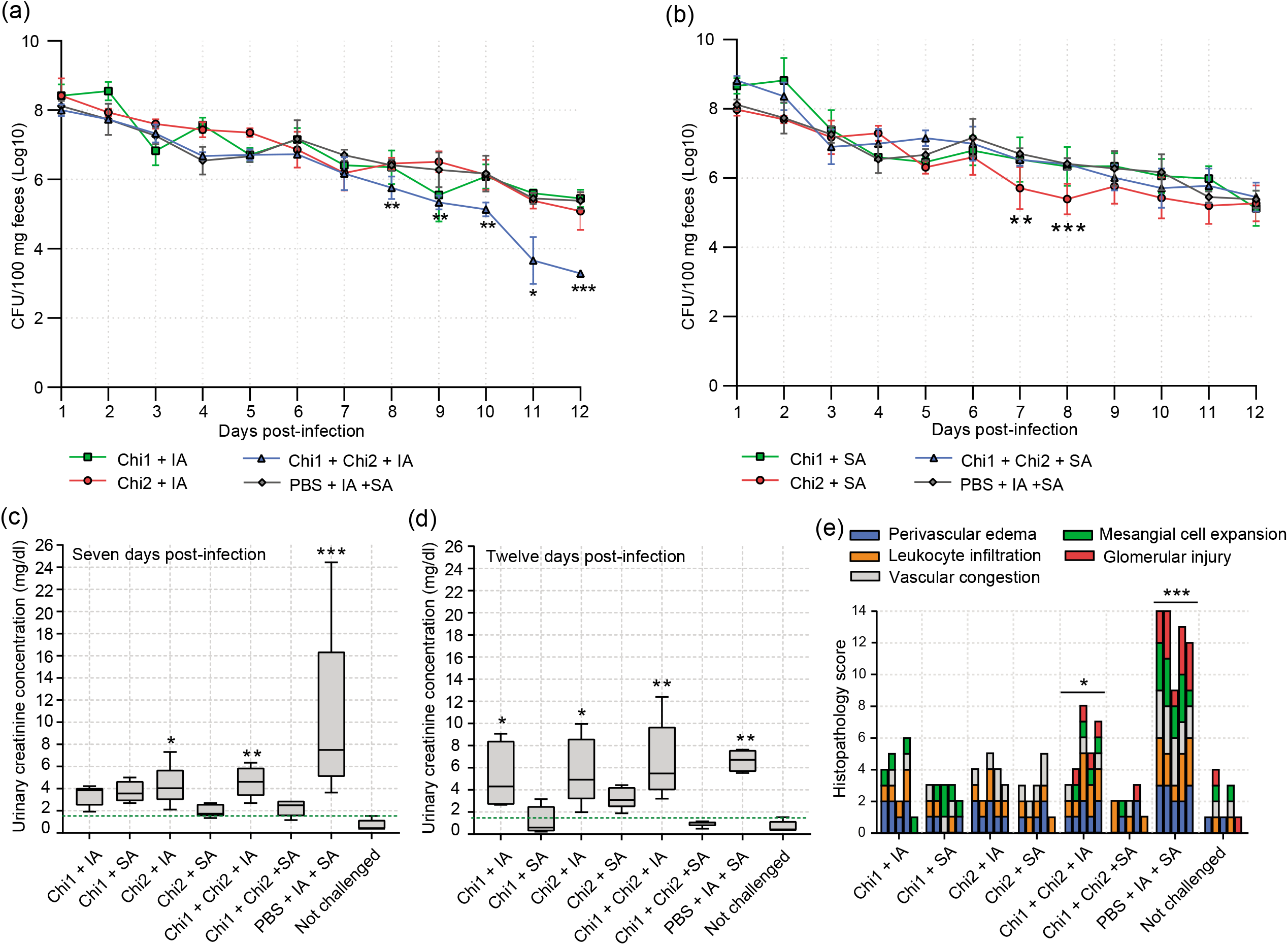
Protection conferred by immunizations with the chimeric antigens. **a and b)** Determination of fecal shedding of STEC O157:H7. Eight mice per group were orally inoculated with 10^9^ CFU of the challenge strain. Fecal pellets were collected daily, weighed, homogenized, and plated on MacConkey agar containing streptomycin. Data are showed as the number of CFU of the challenge strain per 100 mg feces. Error bars represent the standard deviations (s.d.). Differences between immunized mice and the PBS control group were analyzed by a two-way ANOVA with Tukey’s multiple comparison test. Experimental groups are shown as indicated by legend at the bottom. **c and d**) Tukey box plots showing creatinine concentrations (mg/dl) in urine determined from five mice per group on days 7 (c) and 12 (d) post-infection with STEC O91:H21. Differences between experimental groups and uninfected mice were analyzed by Mann-Whitney U test. Dotted green line indicates normal creatinine concentration of 1.5 mg/dl. **e**) Histopathology analysis from kidney tissue obtained from five mice per group on day 12 post-infection with STEC O91:H21. Cellular injuries were classified as not evident, mild, moderate or severe as described in the Methods. Cellular injuries are color coded as indicated in the legend at the top. Differences between experimental groups and uninfected mice were analyzed by a two-way ANOVA followed by Tukey’s multiple comparison test. For all statistical analyses P < 0.0 5 was considered significant.

On day 12 post-infection, mice were euthanized and the level of colonization of the challenge strain in the cecum was determined. Consistent with the above results, recovery of the STEC 86-24 strain was significantly lower in mice immunized with Chi1 plus Chi2 by intramuscular route than in the PBS control group, which corresponded to 2.1 log of protection (**Table 3**). The other experimental groups presented levels of colonization similar to the PBS control group. These data demonstrate that immunization of mice with Chi1 plus Chi2 by intramuscular route induces protective immune responses against intestinal colonization of STEC O157:H7 in this model.

**Table 3.**
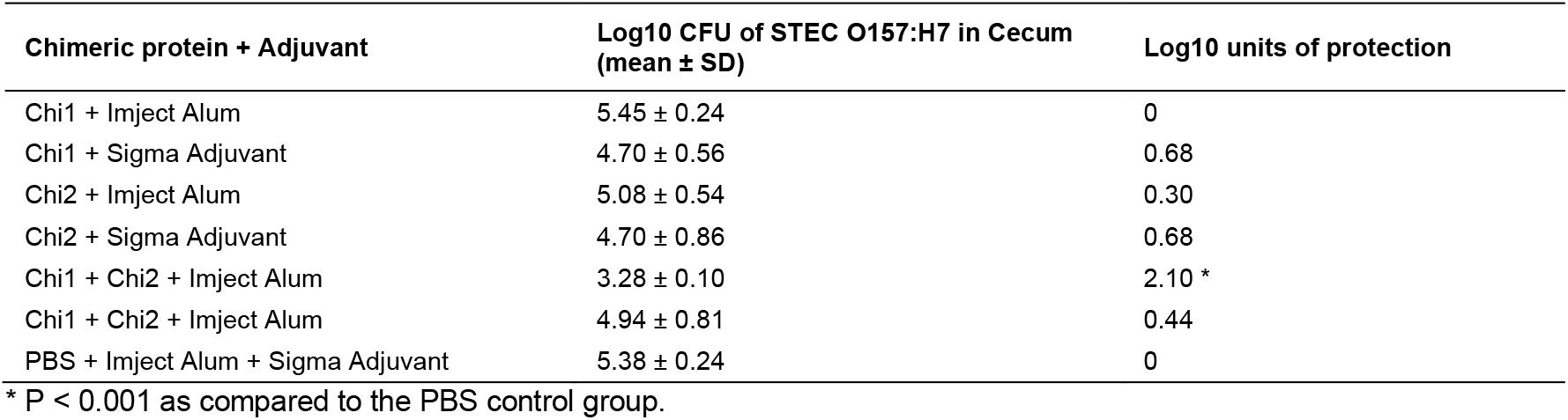
Intestinal colonization of STEC O157:H7 and protection conferred by immunization with chimeric proteins

| Chimeric protein + Adjuvant | Log10 CFU of STEC O157:H7 in Cecum<br>(mean ± SD) | Log10 units of protection |
| --- | --- | --- |
| Chi1 + Imject Alum | 5.45 ± 0.24 | 0 |
| Chi1 + Sigma Adjuvant | 4.70 ± 0.56 | 0.68 |
| Chi2 + Imject Alum | 5.08 ± 0.54 | 0.30 |
| Chi2 + Sigma Adjuvant | 4.70 ± 0.86 | 0.68 |
| Chi1 + Chi2 + Imject Alum | 3.28 ± 0.10 | 2.10 * |
| Chi1 + Chi2 + Imject Alum | 4.94 ± 0.81 | 0.44 |
| PBS + Imject Alum + Sigma Adjuvant | 5.38 ± 0.24 | 0 |

\* P < 0.001 as compared to the PBS control group.

### Chi1 and Chi2 antigens administered alone or in combination by intranasal route avoid renal damage caused by STEC O91:H21

Kidney damage is one of the most severe clinical outcomes that can occur during STEC infection due to the action of Shiga toxins. In streptomycin-treated mice, Stx2d-producing *E. coli* strains have been shown to affect renal function leading to death.^34,35^ For instance, in a previous study we showed that the Stx2d-producing *E. coli* O91:H21 str. V07-4-4 is lethal to mice when they are orally inoculated with a dose of 10^9^ CFU.^10^ However, with a dose lower than 10^5^ CFU of the STEC V07-4-4 strain, mice develop renal pathologies but survive for at least 12 days (unpublished results). Therefore, we conducted further work to investigate whether immunization with the chimeric antigens confers protection against renal damage caused by the STEC V07-4-4 strain. For this, two weeks after the last booster immunization, mice were treated with streptomycin and orally challenged with 10^5^ CFU of the STEC V07-4-4 strain. Our results showed that on days 7 and 12 post-infection, mice intranasally immunized with Chi1, Chi2 or Chi1 plus Chi2 had creatinine levels in urine similar to uninfected mice (**Fig. 5c and 5d**). This was particularly evident in mice immunized with Chi1 plus Chi2. In contrast, mice immunized by intramuscular route and the PBS control group showed significantly higher levels of creatinine in urine than uninfected mice. Moreover, the histopathological analysis of kidney tissue obtained on day 12 post-infection showed that mice immunized by intranasal route had mild or no evident tissue injuries (**Fig 5e**). Conversely, mice immunized by intramuscular route and the PBS control group showed moderate and severe tissue injuries, respectively (**Fig 5e**). Urinary clinical markers such as the number of leukocytes and urobilinogen levels on day 12 post-infection also evidenced the protection achieved by intranasal immunizations (**Table 4**). Together, these results indicate that immunization with the chimeric antigens by intranasal route protects against kidney damage caused by the STEC V07-4-4 strain.

**Table 4.**
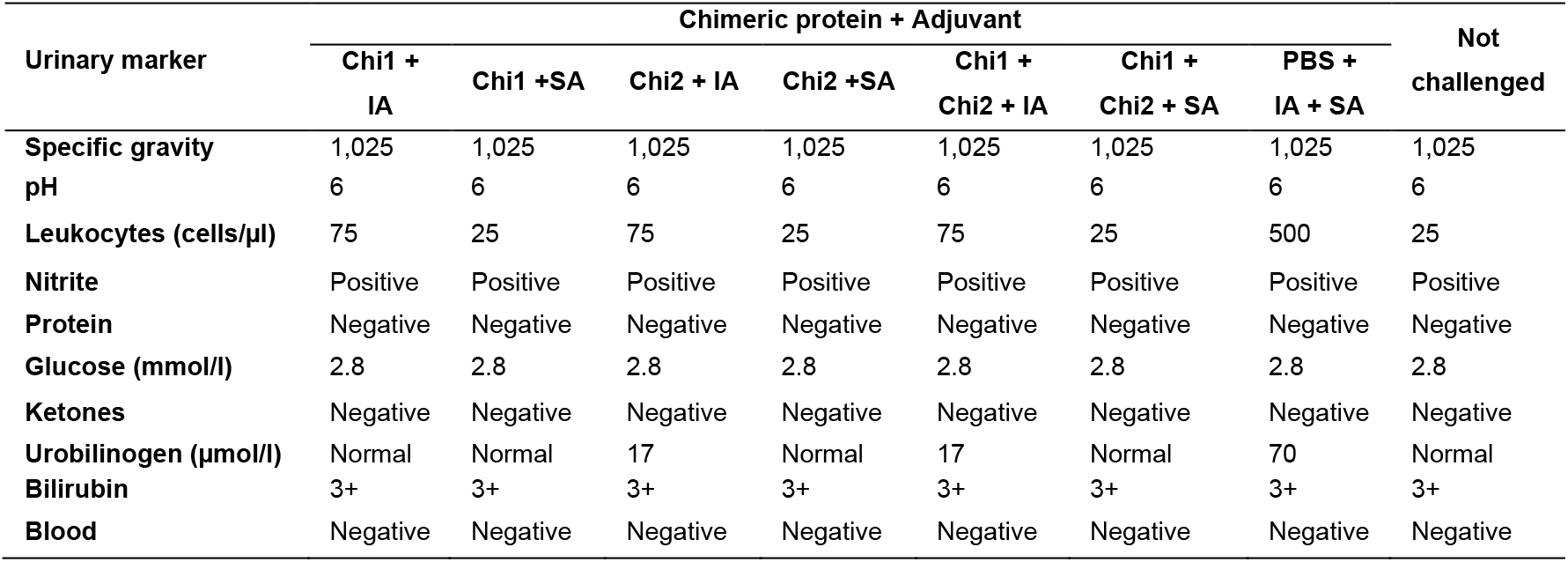
Urinalysis clinical markers assessed at day 12 post-infection

## Discussion

Sera obtained from Chilean-hospitalized pediatric patients diagnosed with HUS, after STEC primoinfection, recognizes antigens such as OmpT, Cah and Hes, as result of a primary immune response to an initial STEC antigen exposure with the development of immunological memory (sera obtained from the convalescent phase with IgG and IgA that recognize STEC antigens). ^13^ Therefore, the key to designing a new vaccine also considers the bacterial target selected for this process. Since STEC infection recurrence is an uncommon process,^36,37^ it is likely that STEC primoinfection can trigger a successful immune memory directed against key bacterial antigens and that remains over time.

Here we demonstrated that immunization of mice with chimeric antigens displaying selected epitopes of OMPs confers protection against intestinal colonization and renal damage caused by STEC. It is well established that a suitable vaccine must be composed of different antigens to boost the immune response with a wide range of protection. Interestingly, the antigens selected for our vaccine design are different from those used in most trials;^62–65^ however, they are widely distributed in STEC and involved in several of its pathogenicity mechanisms.^10,13^ While Cah and Hes are related to the bacterial-host interaction,^9,66^ OmpT participates in the biogenesis of bacterial outer membrane vesicles (OMVs)^58^ and the degradation of antimicrobial peptides like LL-37.^61,67^ To our knowledge, this is the first formulation based on recombinant chimeric proteins that includes a virulence factor exclusive of LEE-negative STEC strains in a vaccine design.

In general, subunit and protein-based vaccines have proven to be safe and to have a defined and homogeneous composition between production batches. This latter aspect is an important advantage over other types of vaccines, which may present complex manufacturing processes, with subsequent regulatory and safety issues.^38–40^ In Gram-negative bacteria, OMPs are primary components interacting with host cells; therefore, vaccines targeting these proteins may be effective by blocking key pathogenic mechanisms.^13,41^ However, as previously mentioned, the purification and efficient production of OMPs pose a major challenge. Thus, construction of water-soluble and stable chimeric antigens displaying epitopes from OMPs is an approach worth exploring.

Our vaccine development approach takes advantage of immunoinformatics tools and *in vitro* assays (epitope mapping, ELISA) to predict and identify linear B-cell epitopes, respectively (**Fig. 1**). Previous studies have also demonstrated the usefulness of immunoinformatics tools for the development of vaccine candidates against STEC and other pathogens.^24,42^ Both, *in silico* and *in vitro* assays, allowed us to select the best epitopes (**Fig. 2, Table 1**,**Table 2**).

Importantly, our results provide proof-in-principle that incorporating selected linear B-cell epitopes into a carrier protein, such as αCah, may result in an increase in antigenicity (**Fig. 3h**). Many autotransporter (AT) proteins and especially their passenger domains are potential vaccine targets.^43,44^ In fact, the pertactin AT from *Bordetella pertussis* is a component of licensed pertussis vaccines.^45^ Thus, other passenger domains from AT proteins could be used as carriers to design chimeric antigens.

The immunization studies showed that the Chi1 and Chi2 antigens, administered alone or in combination, induce humoral responses which remain active until day 42 postimmunization (**Fig 4**). However, the class of antibodies generated was in general dependent on the adjuvant and the administration route. This dependency was most obvious in mice immunized with Chi1 plus Chi2, which elicited significantly higher levels of specific IgG and IgA antibodies in serum when intramuscularly immunized than when intranasally immunized (**Fig. 4a-b**). Further, the sIgA detection in feces after systemic immunization is interesting; however, while this seems to be a controversial issue, there is literature describing this type of results suggesting that vaccines systemically administered can trigger mucosal immune responses.^46^ Many questions await an answer, and one of them is how the immune response moves to the mucosa after systemic immunization, which for some researchers, in addition to producing a paradigm shift, may also mean a modification in vaccine design and delivery.^47^

Secretory IgA is the most abundant immunoglobulin of the mammalian mucosa, playing a fundamental role in the immunity of the gastrointestinal tract.^48^ Therefore, the development of vaccines against intestinal pathogens has traditionally given priority to immunogens that induce significant levels of sIgA. However, since some gastrointestinal infections can be eliminated in the absence of sIgA; other classes of antibodies such as IgM and IgG may also play an important role in the intestinal immunity.^49^ Unfortunately, the presence and effector functions of IgG in the intestinal mucosa have been largely ignored in the literature.^50^ A recent study by Kamada et al., 2015,^51^ revealed that IgG in the murine intestine leads to the selective elimination of a virulent *Citrobacter rodentium* subpopulation by luminal neutrophils. The protective role of IgG against other enteropathogens such as rotavirus has also been demonstrated.^52^

Our challenge experiments using the STEC O157:H7 strain showed that only immunization with Chi1 plus Chi2 by intramuscular route confers protection against intestinal colonization (**Fig. 5a-b, Table 3**). In the murine model of infection, the permanent addition of streptomycin throughout the challenge, promotes STEC O157:H7 colonization by preventing the interference of the microbiota. This situation might explain the decrease in the final part of the protection test (days 8-12), only associated to immune response against STEC O157:H7. Probably, if we had removed the treatment with streptomycin, the interfering activity of the microbiota added to the immune response would have affected early colonization by STEC O157:H7 (before 8 days).^53^ Since intramuscular immunization with Chi1 plus Chi2 did not lead to significant production of specific fecal sIgA antibodies (**Fig. 4d**), it is possible to correlate the protection achieved with other classes of immunoglobulins and more likely with the IgG. On the other hand, although Chi2 includes epitopes of the other two antigens in addition to Cah, it was observed that the set of intranasally immunized mice, yielded a mild immune response against O157:H7 on days 7 and 8, but this was neither sufficiently protective nor maintained over time. Other vaccine candidates that generated significant levels of specific IgG antibodies in serum but not fecal sIgA antibodies have also conferred protection to mice against colonization by STEC O157:H7.^54^ Consequently, our results and those reported by others support the idea of a pivotal role of the IgG antibodies in the defense against enteropathogens. This is a major finding that will be relevant to the development of vaccines against these pathogens by avoiding biases in the selection of the “best” immunogens based mainly on the ability to induce sIgA antibody responses.

Because the challenge studies were performed only to day 12 post-inoculation, it was not possible to determine whether the immune responses triggered by the intramuscular immunization with Chi1 and Chi2 may lead to complete clearance of STEC O157:H7. Long-term protection experiments in mice and other animal models immunized will complement the evaluation of this formulation based on recombinant chimeric proteins.

An effective STEC vaccine may also confer protection against the action of the Stx. STEC export the Shiga toxins along with a number of OMPs and cytoplasmic proteins via outer membrane vesicles (OMVs).^55–57^ These OMVs may be endocytosed in a dynamin-dependent manner by intestinal epithelial cells, and then OMV-associated virulence factors are differentially separated from vesicles during intracellular trafficking. ^57,58^ Recently, it was reported that in the case of Stx2 but not Stx1, once the toxin is internalized, it can be released from eukaryote cells in microvesicles that have exosome markers.^59^ Therefore, immunity against Stx may be mediated by neutralizing Stx-specific antibodies or by immune mechanisms that prevent the toxin from entering the eukaryotic cell.

Our chimeric antigens did not display Stx-associated epitopes. Consequently, the protection conferred by intranasal immunizations against renal damage caused by STEC O91:H21 (**Fig. 5c-e, Table 4**) could be explained by two different mechanisms. The first, the generated immune response might reduce the intestinal colonization of the STEC O91:H21 strain, which could be correlated with a lower release and number of OMVs carrying Stx; however, the intestinal colonization by STEC O91:H21 was not measured. As a result, we cannot conclude that the decrease in colonization correlates with protection against renal damage. The second possible explanation is that sIgA antibodies generated by intranasal immunizations could prevent the release and/or endocytosis of OMVs via immune exclusion. The latter explanation is supported by the fact that OmpT and Ag43 (a Cah homologue protein) are transported in OMVs.^58,60^

In some vaccines, it has been shown that the combination of administration routes, for example mucosal priming followed by systemic boosting or systemic priming followed by mucosal boosting, leads to robust humoral and cellular responses that improve their efficacy.^68,69^ In future studies we will investigate whether the combination of systematic and mucosal immunizations with the Chi1 and Chi2 antigens leads to a more robust and complete immune response characterized by the production of both systemic and secretory antibodies. Also, we will endeavor to reveal the mechanism by which intranasal immunization with Chi1 and Chi2 confers protection against renal damage caused by STEC O91:H21.

Our main focus has always been to protect human health. This is based on the fact that we have seen that the selected antigens are present in a wide range of STEC serotypes previously associated to human illness. However, there are a number of studies that also link these virulence factors to interaction mechanisms between STEC and intestinal epithelial cells in cattle and pigs. In this context, we speculate that our candidate might also be used as a vaccine in animals to prevent STEC colonization, another way to protect the human health.

In conclusion, we developed a promising formulation based on recombinant chimeric proteins that confers protection against STEC intestinal colonization and more relevantly against renal damage caused by Stx. Our study presents interesting results that support the potential use of recombinant chimeras containing epitopes of different antigens of STEC as a preventive strategy.

## Methods

### Bacterial strains and growth conditions

Spontaneously derived streptomycin resistant (Str^r^) mutants of STEC O157:H7 86-24 and STEC O91:H21 V07-4-4 strains were used in this study. Bacterial cultures were routinely grown at 37 °C in Luria-Bertani (LB) broth.

### Human sera

Sera were obtained from 20 pediatric patients in the convalescent phase who presented diarrhea within two weeks prior to HUS diagnosis (HUS sera). Control sera were obtained from two patients with no history of STEC-associated diarrhea. These sera were collected from 1990 to 1993 and from 1999 to 2003 in various healthcare centers in Santiago, Chile, with the written consent of the parents or legal guardians. All procedures were approved by the Ethics Committee of the Facultad de Medicina, Universidad de Chile.

### Peptide Microarray

Epitope mapping assays were performed by PEPperPRINT (Heidelberg, Germany). Briefly, the *ompT, cah* and *hes* sequences were translated into 15, 12 and 10 amino acid peptides with peptide-peptide overlaps of 14, 11 and 9 amino acids. The microarray contained peptides printed in duplicate framed by HA (YPYDVPDYAG) control peptides. Peptide slides were incubated with a mix of three HUS sera at a dilution of 1:100 followed by secondary antibody Goat anti-human IgA (DyLight800) at a dilution of 1:1000, in the presence of the monoclonal anti-HA (12CA5)-DyLight680 control antibody at a dilution of 1:2000. The read-out was performed with a LI-COR Odyssey Imaging System and the image analysis with the PepSlide^®^ Analyzer.

### *In silico* prediction of B-cell and T-cell epitopes

Prediction of B-cell epitopes was done using several tools available at the IEDB server ^70^, including BepiPred 2.0 ^71^, Kolaskar and Tongaonker antigenicity method ^72^ and Ellipro ^73^. Peptides binding to MHC-II molecules were also predicted on the IEDB server ^70^.

### Validation of epitopes by ELISA assay

Seventeen short peptides from 15 to 24 amino acids (**Table 1**) containing linear B-cell epitopes were chemically synthesized at Genic Bio Ltd (Shanghai, China). These short peptides were evaluated for their reactivity to IgG and IgA of individual HUS sera by ELISA. Briefly, 96-well ELISA plates (Nunc Maxisorp or Nunc Immobilizer Amino Plates, ThermoFisher, USA) were incubated with 1.2 μg of each peptide diluted in 100 μl of phosphate-buffered saline (PBS; pH 7.2) overnight at 4 °C. Standard curves were obtained by dilutions in PBS of purified human IgG (Cat. 02-7102, Invitrogen, USA) or IgA (Cat. 3860-1AD-6, Mabtech, USA) ranging from 1.2 μg/ml to 0,0047 μg/ml. Plates were washed three times with PBS containing 0.05% Tween 20 (TPBS) and then incubated with blocking solution (TPBS + 0.5% bovine serum albumin) for 15 min at room temperature (RT). The HUS and control sera were diluted 1:25 (dilution determined from serum titration experiments in a range of 1:10 to 1:100) in blocking solution (100 μl / well) and incubated for 60 min at 37 °C. After six washes with T-PBS, goat anti-human IgG (H + L), peroxidase-labeled (Cat. 04-10-06, KPL, USA) or goat anti-human IgA alpha chain (alkaline phosphatase) (Cat. Ab97212, Abcam, UK), diluted 1:1000 in blocking solution, were added and plates were incubated for 60 min at 37 °C. After six washes with Tris-buffered saline (TBS; pH 7.5) containing 0.05% Tween 20, ABTS^®^ peroxidase substrate (Cat. 50-66-18, KPL, USA) or pNPP substrate (Cat. N2600-10, USBiological, USA) were added and plates were incubated for 12 or 30 min at RT, respectively. The reaction was stopped with 5% sodium dodecyl sulfate or 3 M sodium hydroxide dissolved in distilled water. The absorbance was determined at 405 nm (A_450_) using a Synergy HT microplate reader (Biotek Instruments, USA). Each sample was determined twice in duplicate. The relation between absorbance values and the IgG or IgA concentration of each well was calculated from standard curves using a four-parameter logistic regression in GraphPad Prism 8 software.

### *In silico* modeling and design of chimeric proteins

Predicted three-dimensional structure of Chimera 1 was constructed based on the crystal structures of the Opa60 (PDB_ID: 2MLH)^74^ and OmpT (PDB_ID: 1I78)^75^ proteins. The template for Chimera 2 structure was the crystal structure of the Ag43 protein (PDB_ID: 4KH3).^76^ Comparative modeling of chimeric proteins was performed in Modeller v9 software^77^, using default parameters. The modeled structures were solvated and embedded in a water box using ions (Na+, Cl-) to neutralize the system with the TCL script using VMD software.^78^ The models were optimized with cycles of energy minimization and dynamics using the NAMD 2.12 software.^79^ A molecular dynamics simulation was performed under periodic bordering conditions and isobaric-isothermal set (NPT). The entire system was relaxed by molecular dynamics (MD) simulations using NAMD 2.12 software for 10 ns and subsequently balanced for 30 ns, using the force field CHARMM v2.7.^80^ Quality evaluation and validation of the models were carried out by Ramachandran plot analysis on the RAMPAGE server (http://mordred.bioc.cam.ac.uk/~rapper/rampage.php) and ProSA-web.^81^ Chemical and physical properties of the chimeric proteins were predicted by Protein-Sol^82^ and ProtParam^83^ tools. The modelled structures were visualized with UCSF Chimera 1.10.2.^84^

### Purification of proteins

Synthetic genes and production of Chimera 1, Chimera 2 and αCah proteins were ordered to GenScript (USA). Synthetic genes were optimized for *E. coli* expression and cloned into vector pET30a with N-terminal 6xHis-tag. *E. coli* strain BL21(DE3) was then transformed with recombinant plasmids containing synthetic genes. For purification of recombinant proteins, transformant BL21(DE3) strains were grown in Terrific Broth containing kanamycin (50 μg/ml) at 37° C. When the culture reached an optical density at 600 nm of ~1.2, it was supplemented with IPTG for 4 h. Later, cells were harvested by centrifugation, resuspended with lysis buffer followed by sonication. The sediment obtained after centrifugation was dissolved using urea. Denatured protein was obtained by one-step purification using a Ni-column. The target protein was refolded and sterilized by 0.22 μm filter before being stored in aliquots. The concentration was determined by Bradford protein assay with BSA as standard. The protein purity and molecular weight were determined by standard SDS-PAGE along with Western blot confirmation (Supplementary Figure 1). Target proteins were obtained with purity >85% and endotoxin level <2 EU/mg (LAL Endotoxin Assay Kit, GenScript, Cat. No. L00350). In addition, reactivity of Chimeric and αCah proteins to IgG and IgA of individual HUS was assessed by ELISA assay as described above.

### Immunization studies

All animal experiments were performed at the Universidad de Concepción, Concepción, Chile, following protocols and guidelines approved by the Bioethics Committee of the Faculty of Biological Sciences. Female BALB/c mice (5 to 6 weeks old; purchased from the Instituto de Salud Pública, Santiago, Chile) were randomly distributed into seven experimental groups (each group n=20) and housed in conventional animal facilities with water and food *ad libitum*. Mice were anesthetized with 10 mg/ml of ketamine and 250 μg/ml of acepromazine and immunized by intramuscular (i.m.) or intranasal (i.n.) route with the corresponding protein formulation along with 50 μl of Imject™ Alum Adjuvant (ThermoFisher Scientific, USA) or 20 μl of Sigma Adjuvant System^®^ oil (Sigma-Aldrich, USA), respectively (**Fig. 1d**). The intramuscular immunization was performed in the hamstring muscle and the other group of animals immunized intranasally were anesthetized with a ketamine/xylazine mixture and the corresponding volume of vaccine was administered through the nose. Experimental groups 1,3 and 5 were i.m. immunized with either 20 μg of Chi1, 20 μg of Chi2 or 10 μg of Chi1 plus 10 μg of Chi2, respectively. Experimental groups 2, 4 and 6 were i.n. immunized with either 20 μg of Chi1, 20 μg of Chi2 or 10 μg of Chi1 plus 10 μg of Chi2, respectively. The control group were injected with PBS plus adjuvants. Two booster immunizations were performed on days 15 and 30 using similar amounts of protein formulations and adjuvants. In the BALB/c model, STEC O157:H7 does not cause morbidity or mortality but does allow us to evaluate intestinal colonization. ^85^ In contrast, STEC O91:H21 and its mucus-activated Shiga toxin variant 2d (Stx2d) allowed us to measure kidney damage in infected BALB/c mice.^86,87^

### Sera and feces collection

Sera were obtained from five mice per group by tail vein bleeding on days −2, 14 and 28 before each immunization and at day 42 (two weeks after the last booster immunization), according to conventional techniques. Briefly, blood samples were left at 37 °C for 30 min and then centrifuged at 1,000 x g for 10 min. Supernatant was collected, the complement was inactivated at 56 °C for 30 min and aliquots were stored at −80 °C until IgG, IgA and IgM determinations by ELISA. For secretory IgA (sIgA) measurement, feces were collected on days −2, 14, 28 and 42. Feces were weight, homogenized and diluted to 0.1 g/ml with PBS containing 0.1% sodium azide and 1 mM of phenylmethylsulfonyl fluoride (PMSF). Fecal suspension was centrifuged at 15,000 x g for 5 min at 4 °C, the supernatant fluid recovered and again centrifuged at 15,000 x g for 15 min at 4 °C and stored at −80 °C until use.

### Measurement of humoral response

Chimeric proteins were diluted to 1 μg/ml in carbonate buffer (pH 9.6) and used to coat polystyrene 96-well high-binding ELISA plates (100 μl/well; Nunc-Immuno plate with MaxiSorp surface). After overnight incubation at 4 °C, plates were washed with washing buffer (Tris-buffered saline [pH 7.4] with 0.05% Tween 20) and blocked with 0.8% gelatin in TPBS for 1 h at 37 °C and then incubated with either sera or supernatant from fecal suspensions, at a dilution of 1:100, for 2.5 h at room temperature and washed four times. Next, isotype-specific goat anti-mouse HRP conjugates (BioLegend, USA) were added (100 μl/well) at a dilution of 1:1000 and incubated for 1 h min at room temperature followed by washing with TPBS. Then, 200 μl/well of OPD Peroxidase Substrate (Cat. P9187-5SET, Sigma-Aldric, USA) was added for 30 min. The reaction was stopped with 50 μl/well of 2 N H_2_SO_4_ and the absorbance at 450 nm was measured on a microplate reader.

### Challenge studies

Two weeks after the last booster immunization (day 45), the infection experiments were performed in the streptomycin-treated mouse model of STEC infection as described elsewhere^87,88^ with minor modifications. Briefly, mice were given water *ad libitum* containing streptomycin (5 g/l) 24–48 h prior to inoculation and for the duration of the experiment. Feces were documented to be free of streptomycin-resistant *E. coli* at the time of inoculation. STEC O157:H7 86-24 and STEC O91:H21 V07-4-4 strains were grown overnight in agitated LB broth containing 50 μg/ml streptomycin at 37 °C. Cultures were centrifuged, washed once with PBS and resuspended in a 20% sucrose (w/v) and 10% NaHCO3 (w/v) solution in sterile water to 1 × 10^10^ CFU/ml (STEC 86-24 strain) or 1 × 10^6^ CFU/ml (STEC V07-4-4 strain). Prior to inoculation, mice were starved of food and water overnight (12 h). The next morning mice were orally infected by pipette feeding with 100 μl of bacterial suspension containing 10^9^ CFU or 10^5^ CFU of STEC 86-24 or STEC V07-4-4 strains, respectively. After challenge, food and water were reintroduced and provided *ad libitum*. The fecal shedding of the 86-24 strain was recorded daily for 12 days. For this, feces were collected, weighed, homogenized, suspended in 1 ml PBS and, after serial dilutions, plated on MacConkey agar plates supplemented with streptomycin (50 μg/ml) for bacterial counts. For determination of intestinal colonization of the STEC 86-24 strain, 12 days after challenge (day 57), mice were euthanized, and cecum was collected under aseptic conditions, homogenized and diluted in PBS. Suspensions were serially diluted and plated on MacConkey supplemented with streptomycin (50 μg/ml) for bacterial counts. Log_10_ units of protection were obtained by subtracting the mean Log_10_ CFU for each experimental group from the mean Log_10_ CFU of the PBS control group. Mice inoculated with the STEC V07-4-4 strain were used to measure kidney damage (*vide infra*).

### Urinalysis

Urine samples were collected as previously described ^89^, on days 45, 51 and 57 (0, 7 and 12 post-infection, respectively) from mice infected with the STEC V07-4-4 strain. Biochemical estimation of urine creatinine concentration was assessed using the Creatinine Kit (BioSystems, Spain) according to the manufacturer’s instructions. Other clinical urine markers were measured by using Combur10 Test®M semiquantitative test strips (Roche Diagnostics GmbH, Germany). Each test strip consists of colorimetric reaction spots for 10 markers: specific gravity (1.000 to 1.030), pH (5.0 to 9.0), leukocytes (range, negative to 500 cells/μL), nitrites (negative or positive), proteins (negative to 500 mg/dl), glucose (negative to 55 mmol/l), ketones (negative to 15 mmol/L), urobilinogen (normal to 200 μmol/L), bilirubin (negative to +3) and blood (negative; trace of non-hemolyzed; or hemolyzed, 10 to 250 cells/μL). Each square was wet with a drop of urine and the marker value was determined through comparison with a colorimetric standard.

### Histopathological analysis of kidney tissue

For the histological analysis of kidney tissue, mice infected with the STEC V07-4-4 strain were euthanized at day 57, the kidneys collected, fixed in 10% formaldehyde (pH 6.9), embedded in paraffin wax for sectioning at 5 μm and stained with hematoxylin and eosin (H/E). Pathological evaluation of H/E-stained tissue sections was carried out by a pathologist blinded to the experimental design. Histopathological changes were evaluated by the degree of perivascular edema, leukocyte infiltration, vascular congestion, mesangial cell expansion and injury of the glomerular filtration barrier (glomerular hypertrophy or glomerular hypoperfusion). Each sample was quantitated by ten randomly selected fields with the following criteria: 0, no damage; 1, <25%; 2, 25–50%; 3, >50%; 4, >75% of damage. Differences between experimental groups were evaluated by a one-way ANOVA followed by Tukey’s multiple comparisons test.

## Data availability

The data that support the findings of this study are available on request from the corresponding author R.V. Amino acid sequences of Chimeric proteins and identified epitopes are not publicly available due to legal restrictions and an ongoing international patent application.

## Acknowledgments

This study was supported by FONDEF ID16I10140 and FONDECYT 1161161 grants. We thank Dr. Helen Lowry for the careful revision and edition of the manuscript. We thank Dr. Mauricio Farfán for sharing the STEC O157:H7 str. 86-24.

## Author contributions

Conceptualization and experimental design: Roberto Vidal, David A. Montero, Felipe Del Canto, Juan C. Salazar and Angel Oñate. Data acquisition: Sandra Céspedes, Leandro Cádiz, José Reyes, Mauricio Arenas and David A. Montero. Data analysis and interpretation: Roberto Vidal, David A. Montero, Felipe Del Canto, Juan C. Salazar and Angel Oñate. Writing – original draft: David A. Montero; Review and edition: Roberto Vidal, Felipe Del Canto, Juan C. Salazar, Mauricio Arenas and Angel Oñate.

